# Hepatic ketogenic insufficiency blunts exercise-induced energy expenditure and alters mitochondrial proteins in skeletal muscle

**DOI:** 10.1101/2025.09.16.676620

**Authors:** Xin C Davis, Colin S McCoin, Sebastian F Salathe, Edziu Franczak, Julie A Allen, Eric D Queathem, Kyle L Fulghum, Patrycja Puchalska, Peter A Crawford, John P Thyfault, E. Matthew Morris

## Abstract

Ketone body (KB) utilization increases during fasting and exercise due to enhanced hepatic fatty acid oxidation and KB production via the rate-limiting mitochondrial enzyme hydroxymethylglutaryl-CoA synthase (HMGCS2). Since KB metabolism intersects with multiple metabolic pathways, and skeletal muscle KB catabolism rises during exercise, we tested the hypothesis that liver-specific HMGCS2 knockouts (KO) would have reduced energy expenditure (EE) and changes in the mitochondrial proteome of skeletal muscle with chronic exercise through voluntary wheel running (VWR), time-restricted feeding (TRF), or both combined to boost hepatic KB production and utilization. Control (CON) and HMGCS2 knockout (KO) mice (n=6-8 per group) underwent sedentary ad libitum feeding (SED+AL), SED+TRF, VWR+AL, and VWR+TRF for 16 weeks, with whole-body EE measured using indirect calorimetry. In CON mice, VWR increased total EE by 19.5% and non-resting EE by 50% under AL conditions, and total EE by 16% and non-resting EE by 47.9% under TRF conditions. However, the EE increases seen with VWR did not occur in KO mice. Proteomic analysis revealed that the loss of liver HMGCS2 significantly impacted proteins involved in metabolic processes within skeletal muscle, including reduced oxidative phosphorylation (OXPHOS) protein expression in SED KO mice compared to sedentary CON. Notably, VWR restored OXPHOS protein expression in the muscle of the liver HMGCS2 KO but did not alter it in the CON. Furthermore, muscle from liver HMGCS2 KO mice had elevated expression glycolytic pathways in sedentary and VWR conditions. These results indicate that hepatic ketogenic deficiency (HMGCS2 KO) diminishes exercise-induced increases in EE and uniquely impacts baseline and exercise-related adaptations in the metabolic and mitochondrial proteome of skeletal muscle.

## Introduction

Over the past several decades, interventions designed to increase serum ketone bodies (KBs) have been purported to increase fat burning and reduce hunger. Fasting, exercise, and/or ketogenic diets (high fat/low carbohydrate) are commonly employed strategies to stimulate ketogenesis (1). During states of low carbohydrate availability, hepatic fatty acid oxidation is elevated to increase the rate of acetyl-CoA formation, which serves as a precursor for acetoacetate (AcAc) via the rate-limiting enzyme: 3-hydroxymethylglutary-CoA synthase (HMGCS2). AcAc is further reduced to β-hydroxybutyrate (β-OHB). Liver-derived KBs serve as additional oxidative fuel sources for several prominent metabolic tissues like the brain, heart, and skeletal muscle (2, 3). Beyond serving as metabolic substrates under fasting or carbohydrate-deficient conditions, KBs also attenuate peripheral glucose utilization, induce anti-lipolytic effects on adipose tissue, decrease proteolysis in skeletal muscle, and impact whole body energy expenditure (EE) (4, 5). Several studies have shown that increasing KB levels protects against oxidative stress and muscle atrophy while elevating oxidative metabolism (6). A recent review highlighted the mixed results of altering circulating KB levels with different diet or supplement strategies (low carbohydrate, medium chain triglycerides, and intake of ketone esters) on energy balance, concurring that tightly controlled mechanistic studies are needed to better understand the effects of KB on whole body energy metabolism (7). Despite various studies suggesting the importance of KBs in energy metabolism, the impact of impaired ketogenesis on whole-body and skeletal muscle energy metabolism remains relatively unknown, especially during physiological stimuli like fasting and exercise.

Skeletal muscle can rapidly remodel metabolic pathways to meet metabolic demands and is the primary site for exercise-induced increases in EE as well as a primary regulator of resting EE. Acute exercise increases skeletal muscle KB utilization (8) and total capacity to oxidize KB in muscle is increased with chronic exercise training (9). Elevated KB levels have also been shown to induce alterations in both the skeletal muscle transcriptome (10) and proteome (11). Despite these findings, little is known regarding whether decreased systemic KB availability affects the skeletal muscle proteome in a sedentary condition or in response to chronic physiological stimuli that induce ketogenesis. Being one of the main KB consumers and a source of metabolic intermediates, muscle can significantly affect whole-body energy metabolism during exercise. Thus, how impaired hepatic ketone production impacts skeletal muscle in sedentary and exercise conditions remains unknown.

Therefore, the first objective of this study was to investigate whether insufficient hepatic ketogenesis would affect whole-body EE in mice undergoing long-term interventions known to induce ketogenesis. Time-restricted feeding (TRF) and exercise increase hepatic ketogenesis, and the combination of both produces a synergetic effect that further elevates ketogenesis and whole body KB turnover (8). Thus, we measured total, resting, and non-resting EE via indirect calorimetry in both control (CON) and hepatic-specific HMGCS2 knockout (KO) mice undergoing chronic TRF, exercise via voluntary wheel running (VWR), or TRF+VWR. Secondly, we investigated if hepatic ketogenesis deficiency and subsequent compromised ketone utilization would alter exercise-induced proteome adaptation in gastrocnemius muscle. Our findings revealed that impaired ketogenesis significantly blunts the capacity of voluntary exercise to increase total and non-resting EE while also changing the skeletal muscle proteome under sedentary and chronic exercise conditions.

## Methods

### Ethical approval and experimental protocol

Liver-specific HMGCS2 knockout mice (KO) were generated via a cross of the Albumin-Cre strain (Jackson Laboratories, strain #035593) to the floxed Hmgcs2 allele on a C57BL/6NJ background as described previously (12). Male C57BL/6NJ control (CON) and liver-specific HMGCS2 KO mice, (n=6-8 per treatment group) were individually housed at 22 °C on a reverse light cycle (Dark: 10:00-22:00) with *ad libitum* access to water and were allocated to one of four treatments at average 34-35 weeks of age. Treatments included AL, TRF, VWR, or VWR+TRF. The following sample size was included in each group (SED+AL; n= 6 for controls, n= 8 for KO), (TRF+SED; n=7 for controls, n=8 for KO), (VWR+AL; n=6 for controls, n=8 for KO), and (VWR+TRF; n=7 for controls and n=8 for KO). The AL group had *ad libitum* access to low fat diet [LFD; D12110704: 10% kcal fat, 70% carbohydrate (3.5% kcal sucrose), and 20% kcal protein with an energy density of 3.85 kcal/g, Research Diets, New Brunswick, NJ, USA] while the TRF group were fasted from 8:30-16:30 Monday through Friday, with *ad libitum* access to food on Saturday and Sunday. Food was pulled 90 minutes before the dark cycle (08:30) to preclude food consumption occurring prior to the start of the dark cycle. In addition, mice regularly start running at the onset of the dark cycle, thus VWR+TRF induced a treatment that is on par with “fasted exercise,” resulting in a synergetic ketogenesis response before food was replaced at 16:30. The mice had *ad libitum* access to food for the remaining 5.5 hours of the dark period (from 16:30 to 22:00). The VWR group had *ad libitum* access to vertical running wheels (ENV-047V, Med Associates Inc) and running distance was continuously recorded throughout intervention. The interventions occurred for 16 weeks before the mice were placed in indirect calorimetry cages for 4 days. All animal use was in accordance with the National Institutes of Health (NIH) *Guide for the care and use of laboratory animal* and conformed to the principles specified in protocols approved by the University of Kansas Medical Center and University of Minnesota Institutional Animal Care and Use Committees (IACUCs).

### Anthropometric

Body weight and body composition were collected weekly as well as prior and after indirect calorimetry. Food intake was measured weekly during the 16-week intervention and during indirect calorimetry. Body composition was determined via MRI (EchoMRI-1100, EchoMRI, Houston, TX), with the fat-free mass calculated as the difference between body mass and fat mass.

### Serum Ketone measurements

Serum D-beta-hydroxybutyrate (D-βOHB) and acetoacetate (AcAc) concentrations were quantified at the time of tissue collection after a 2 hour fast via UHPLC-MS/MS as reported previously (13). Briefly, AcAc and D-βOHB were extracted from 10 µL of serum into 40 µL of ice cold (−20°C) methanol: acetonitrile (1:1) spiked with [U-^13^C4]AcAc and D-[3,4,4,4-^2^H2]βOHB internal standards at a final concentration of 50 µM. Samples were spun at 18k x g for 10 min at 4°C then 1 µL of the supernatant was injected onto a C_18_ Cortecs UPLC T3 column (150 x 2.1, 1.6 µm) (Waters 186008500), separated via reverse-phase chromatography, and detected via parallel reaction monitoring (PRM) on a QExactive Plus hybrid quadrupole-orbitrap mass spectrometer, equipped with a heated electrospray ionization (ESI) source operated in negative ionization mode.

### Indirect Calorimetry

Energy metabolism was measured using Promethion indirect calorimetry metabolic monitoring system, as previously described (Sable Systems International, Las Vegas, NV) (14, 15). Food restriction for TRF groups was continued within the indirect calorimetry system, and all VWR groups-maintained access to wheels while in the system. Mice were acclimated in the indirect system cages for 3 days, with data collection occurring on the 4^th^ day. Respiratory exchange rate (RER) was calculated as V_CO2_/V_O2_. Total EE was calculated as the mean EE per hr using a modified Weir equation [EE (kcal/hr) = 60∗ (0.003941 × V_O2_+0.001106 × V_CO2_) and extrapolated to 24 hrs. Resting EE was calculated from the 30-minute period with the lowest EE and extrapolated for a 24-hour period. Non-resting EE was calculated as total EE minus resting EE. Ambulatory cage activity was quantified by total X, Y, and Z beam breaks, as previously described (14). For VWR groups, ambulatory cage activity was added to wheel meters during data collection to determine Total Meters. ‘Fasting’ and ‘fed’ RER, EE, and Total Meters were calculated by averaging the data during the 8-hr fasting period (8:30 to 16:30) and the 16-hr fed period separately. Efficiency of movement was determined from the following equation: non-resting EE/Cage activity (Fletcher & MacIntosh, 2017). Linear mixed-effects modeling using a three-way ANCOVA was performed as previously described (see below) (14).

### Tissue collection

Food was pulled 2 hours prior to tissue collection at 10:00 am. Mice were anesthetized with an intraperitoneal injection of 25∼30 μl of 50 mg/ml phenobarbital followed by exsanguination. Blood was collected via cardiac puncture. Gastrocnemius muscle was dissected, frozen in liquid nitrogen, and stored at -80°C and processed for proteomics analysis.

### Gastrocnemius muscle preparation for LC-MS/MS analysis

Proteins were extracted by adding 30ul per mg of tissue RIPA buffer containing protease inhibitors and incubating on ice for 30 minutes followed by sonicating in a water bath for 15 minutes. A Bicinchoninic Acid assay (BCA) assay was performed to determine protein content and based on the concentrations, 50 µg of protein from each homogenized and clarified mouse gastric sample was transferred to a new tube and volumes were increased to 50µl using 50 mM TEAB. Samples were reduced with 5 mM TCEP followed by incubation at 55 °C for 30 minutes. Cysteines were alkylated by the addition of 375 mM iodoacetic acid (IAA) to a final concentration of 10 mM followed by incubation at room temperature in the dark for 30 minutes. Ice cold acetone was added at a 1:5 ratio followed by incubation at -20 °C overnight. After precipitation, samples were centrifuged at 14,000 x g at 4 °C for 10 minutes to pellet the proteins. The supernatant was removed, and proteins were allowed to air dry on the bench top for 15 minutes. The proteins were resuspended in 100µl of 50 mM TEAB (pH 8) with 2 mM CaCl2 and proteins were digested by adding 500ng of trypsin and incubating overnight at 37 °C with shaking at 500 RPM (Thermomixer, Eppendorf). Digestion reactions were quenched by the addition of 10% formic acid to a final concentration of 1%. Digested samples were centrifuged at 10,000 x g for 10 minutes to remove particulates and supernatant was transferred to a fresh tube. Peptide concentrations were measured using a Nanodrop spectrophotometer (Thermo Scientific) at 205 nm.

### Proteomic LC-MS/MS detection and data analysis

Samples were injected using the Vanquish Neo (Thermo) nano-UPLC onto a C18 trap column (0.3 mm x 5 mm, 5 µm C18) using pressure loading. Peptides were eluted onto the separation column (PepMap™ Neo, 75 µm x 150 mm, 2 µm C18 particle size, Thermo) prior to elution directly to the MS. Briefly, peptides were loaded and washed for 5 minutes at a flow rate of 0.350 µL/min at 2% B (mobile phase A: 0.1% formic acid in water, mobile phase B: 80% ACN, 0.1% formic acid in water). Peptides were eluted over 100 minutes from 2-25% mobile phase B before ramping to 40% B in 20 min. The column was washed for 15 min at 100% B before re-equilibrating at 2% B for the next injection. The nano-LC was directly interfaced with the Orbitrap Ascend Tribrid MS (Thermo) using a silica emitter (20 µm i.d., 10 cm) equipped with a high field asymmetric ion mobility spectrometry (FAIMS) source. The data were collected by data dependent acquisition with the intact peptide detected in the Orbitrap at 120,000 resolving power from 375-1500 m/z. Peptides from samples with a charge of +2-7 were selected for fragmentation by higher energy collision dissociation (HCD) at 28% NCE and were detected in the ion trap using rapid scan rate. Dynamic exclusion was set to 60s after one instance. The mass list was shared between the FAIMS compensation voltages. FAIMS voltages were set at -45 (1.4 s), -60 (1 s), - 75 (0.6 s) CV for a total duty cycle time of 3s. Source ionization was set at 1700 V with the ion transfer tube temperature of 305 °C. Raw files were searched against the mouse protein database downloaded from Uniprot on 09-26-2023 using SEQUEST in Proteome Discoverer 3.0 (16).

### Gastrocnemius muscle proteomics data analysis

All proteins identified from proteomics analysis were included for pathway analysis using Ingenuity Pathway Analysis (IPA, Qiagen). log2FC and p-value were used to identify up- and down-regulated cellular processes, as previously performed (17). Mitocarta3.0 (18) was used to examine mitochondrial-specific pathways identified as enriched within the proteomics dataset, as done previously. (17, 19). Briefly, MS2 raw intensity of all mitochondrial proteins were summed and corrected for the total raw intensity of the entire proteome to compare protein abundance differences across all groups.

### Statistics Analysis

For each group, data are presented as mean ± standard error mean (SEM). Outliers were identified and removed using the two-standard-deviations test. A three-way ANOVA was used to test the effects of genotype (CON vs. KO), exercise (VWR vs. SED), and feeding status (TRF vs. AL) for anthropometric and indirect calorimetry data (Prism 10, GraphPad). Two-way ANOVA was explicitly used when assessing wheel meters in the VWR groups. Additionally, three-way ANCOVA was used to test if differences in fat mass and fat-free mass were significant covariates of total EE, non-resting EE, and resting EE. Adjusted marginal means and effect size (partial eta^2^) based on differences in body composition were also calculated (SPSS Statistics, IBM Corp., Armonk, NY, USA). For analysis of protein content within the proteomics data, two-way ANOVA was conducted (genotype [control vs. KO] vs. interventions [SED+AL, VWR+AL, and VWR+TRF]). When significant interactions and/or main effects were detected, Fisher’s LSD *post hoc* analysis was performed. Main effects were only described when all within-group *post hoc* comparisons were significant.

## Results

### Anthropometrics and ketosis

There was no significant difference in final body weight between CON and KO mice, or with VWR or TRF (Table 1). TRF decreased fat mass in both CON and KO compared to SED groups. Fat-free mass was reduced by ∼8% in KO compared to CON within SED + AD LIB. Fat-free mass also increased by ∼9% in KO VWR group compared to KO in SED fed AD LIB (Table 1, p<0.05). VWR increased average weekly food intake and energy intake (∼28%) (p<0.05) compared to SED in both CON and KO. After a 24 hr fast, blood KB levels were elevated in CON TRF mice compared to CON AD LIB mice (Table 1, p<0.05), with no effect of VWR. However, as expected, KO mice showed no increase in blood KB induced by TRF or VWR. In the 2h-fasted state (when ketone bodies circulate at lower concentrations), application of our sensitive UPLC-MS/MS method showed that KO mice showed consistently lower KB concentrations than controls across all conditions (Supplemental Figure 1), with no statistically significant effects of the TRF or VWR interventions. The average VWR distance per day did not differ between TRF and AD LIB, or by genotype, over the course of VWR exposure (Table 1).

**Table 1.**

|  | SED |  |  |  | VWR |  |  |  |
| --- | --- | --- | --- | --- | --- | --- | --- | --- |
|  | AD LIB |  | TRF |  | AD LIB |  | TRF |  |
|  | CON | KO | CON | KO | CON | KO | CON | KO |
| **BW (grams)** | 38.5 ± 1.4 | 35.4 ± 1.9 | 37.3 ± 1.7 | 36.5 ± 1.1 | 35.4 ± 2.1 | 36.7 ± 1.5 | 36.7 ± 2.5 | 34.1 ± 1.3 |
| **FM (grams)** | 11.2 ± 0.9 | 10.6 ± 1.5 | 10.9 ± 1.1 | 10.9 ± 0.8 | 8.5 ± 1.6 | 9.0 ± 1.6 | 9.1 ± 1.8 | 8.6 ± 1.2 |
| **FFM (grams)** | 25.6 ± 0.4 | 23.5 ± 0.3<sup>++</sup> | 24.6 ± 0.4 | 24.4 ± 0.4 | 26.0 ± 0.8 | 25.6 ± 0.6<sup>**</sup> | 25.9 ± 0.8 | 25.1 ± 0.5 |
| **Weekly Energy intake (kcal)** | 83.6 ± 1.30 | 78.7 ± 2.1 | 80.2 ± 3.2 | 79.3 ± 1.3 | 102.0 ± 2.8* | 103.9 ± 1.9* | 101.9 ± 2.5* | 104.1 ± 2.6* |
| **Wheel meter per day (km)** | - | - | - | - | 4.0 ± 0.7 | 3.2 ± 0.9 | 3.5 ± 0.7 | 5.0 ± 1.0 |

Data are reported as the estimated mean ± SEM, n=6-8/group. \* main effect of VWR p<0.05. && treatment within genotype and activity, ++ genotype within activity and treatment p<0.05, \*\* Activity within genotype and treatment p<0.05.

### Male HMGCS2 KO mice display lower VWR-induced energy expenditure

Indirect calorimetry data were collected over 24 hours, and was performed while mice were maintained through their respective daily interventions of VWR or SED and the 8-hour food restriction (TRF) or AD LIB feeding. VWR increased total EE in CON (11%) mice compared to the SED (Figure 1A, p<0.05). Interestingly, this increase did not occur in KO VWR mice, suggesting that hepatic ketogenic deficiency blunts exercise-induced total EE. Quantification of the two main components of total EE: resting and non-resting EE, revealed no differences in resting EE between genotypes, activity, or treatment groups (Figure 1B). In contrast, CON VWR had ∼40% greater non-resting EE compared to CON SED (Figure 1C, p<0.05), while VWR did not increase non-resting EE in KO mice compared to SED (Figure 1C).

**Figure 1.**
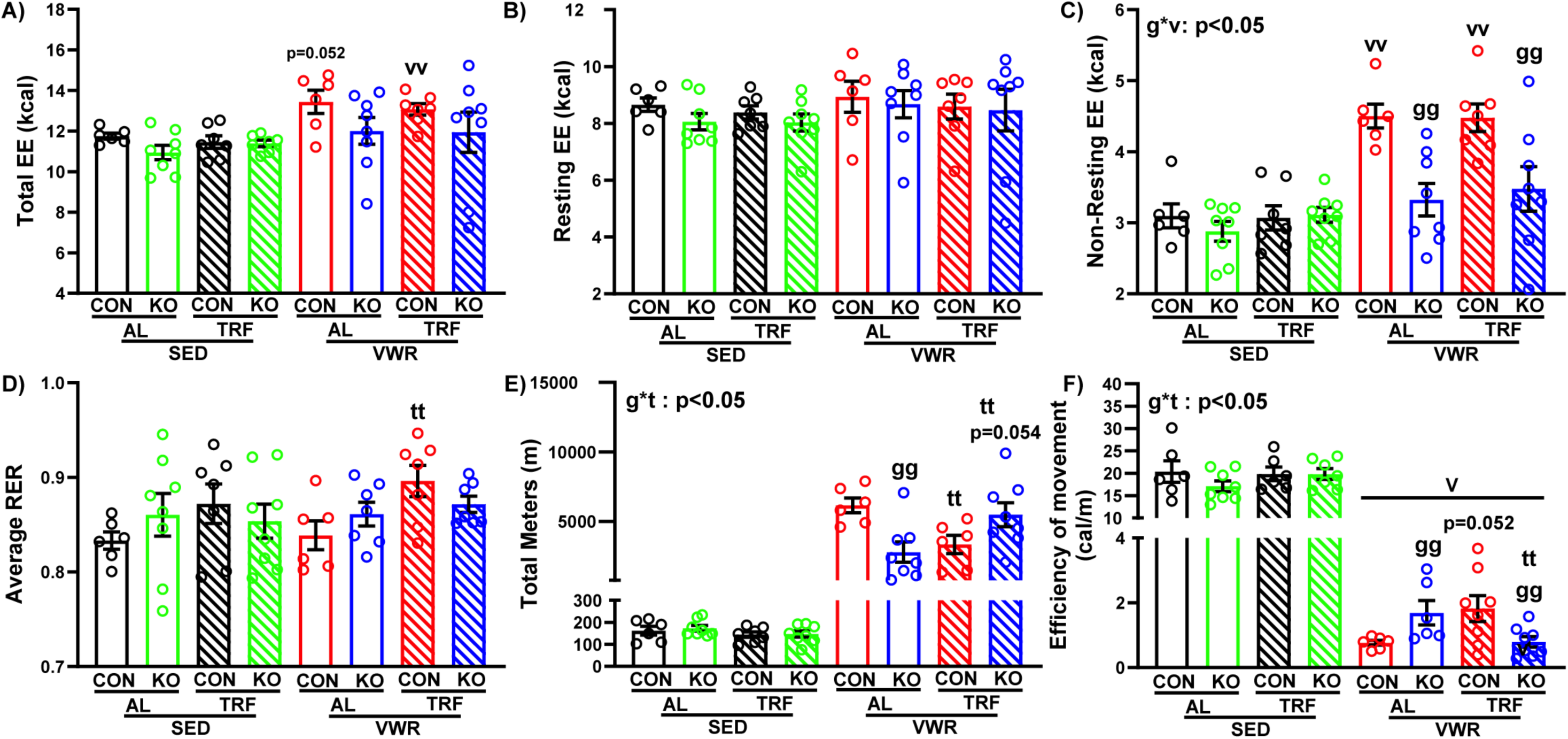
HMGCS2 KO display lower EE phenotype. (A) Total energy expenditure (EE); (B) Resting energy expenditure (REE); (C) Non-Resting energy expenditure (Non-REE); (D) Average respiratory exchange ratio (RER); (E) Total Meters (all meters of ambulatory movement and wheel running) (F) Efficiency of movement defined as Non-REE/All meters Data presented as mean +/- SEM (n=6-8 per group). g* t: genotype*TRF interaction p<0.05, v: p<0.05 main effect of VWR, gg: p<0.05 genotype within feeding and activity, tt: p<0.05 feeding within genotype and activity, vv: p<0.05 activity within genotype and feeding.

Exercise and TRF impact substrate utilization by increasing systemic fat metabolism and enhancing ketogenesis (8). We hypothesized that TRF would lead to greater fat utilization and, consequently, a lower RER in the CON TRF groups. Contrary to our expectations, TRF in CON SED mice did not change RER compared to AD LIB, while CON VWR TRF mice had a higher 24 hr average RER compared to AD LIB (Figure 1D, p<0.05), averaged over an entire 24-hour period. RER was not different between KO mice and CON mice; moreover, TRF and VWR did not influence RER in the KO mice quantified over an entire 24-hour period (Figure 1D). EE is highly correlated with body mass and fat-free mass (20) and is impacted by adiposity (21, 22). To assess whether differences in EE were due to the differences in body composition, we performed an analysis of covariance using fat mass and fat-free mass as covariates to generate estimated marginal means and assess covariate effect size (Table 2). While both fat and fat-free mass were significant predictors of variability in total and resting EE (p<0.05), the effect sizes for fat and fat-free mass were small, and the comparisons of the estimated marginal means were not different than those in Figure 1.

**Table 2.**
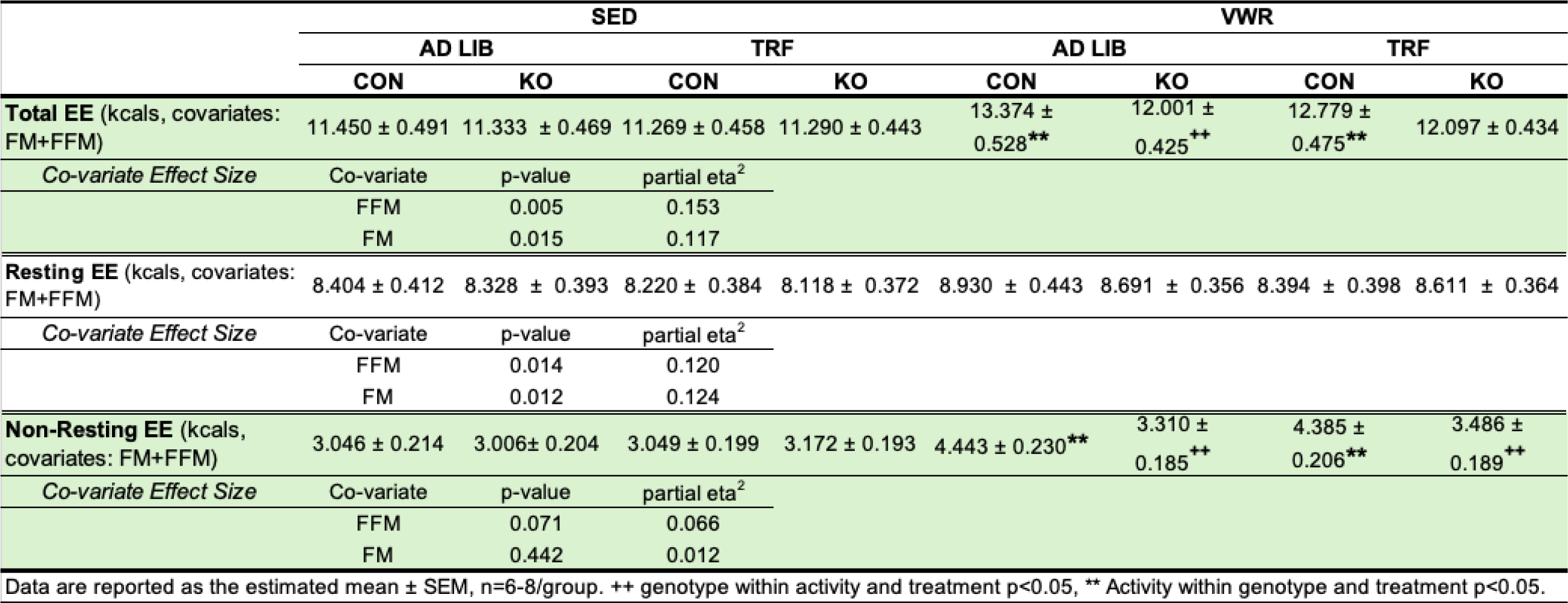

We also examined whether genotype, VWR, or TRF influenced total movement in the indirect calorimetry cages. Total Meters encompasses both ambulatory cage movement and VWR distance over a 24-hour period in the calorimetry cages. Total Meters were not affected by genotype or treatment in SED mice (Figure 1E), whereas VWR significantly increased cage activity across all groups as expected (p<0.05). Within the VWR groups, an interaction between genotype and TRF was observed (Figure 1E, p<0.05). Total Meters was lower in CON TRF mice compared to CON AD LIB (p<0.05). However, while KO AD LIB mice showed decreased Total Meters than CON KO AD LIB mice (p<0.05), KO TRF mice exhibited greater Total Meters than KO AD LIB (p<0.05) and Total Meters of KO TRF trended higher than CON TRF (p=0.054). To explore whether this interaction was due to differences in the ambulatory cage movement versus VWR components of cage activity, we compared wheel meters (Figure 3A) and ambulatory cage activity in VWR groups (Figure 3B). The interaction between genotype and TRF was again evident in wheel running distance, but not in ambulatory cage activity. Given the differences in non-resting EE and cage activity between genotypes and treatments, we calculated movement efficiency, defined as non-resting EE divided by cage activity (cal/meter). A higher value indicates less efficiency, and a lower value indicates greater efficiency (23). As expected, VWR significantly increased movement efficiency in both genotypes (Figure 1F, p<0.05). An interaction between genotype and treatment was also observed in VWR mice (p<0.05). CON TRF mice tended to be less efficient than AD LIB (p=0.052). While KO AD LIB mice demonstrated lower efficiency than CON (p<0.05), KO TRF mice showed greater efficiency than both KO AD LIB and CON TRF (p<0.05). Overall, these findings indicate that during increased activity (wheel running), hepatic ketogenic insufficiency reduces EE and interacts with food restriction to influence movement efficiency.

**Figure 2.**
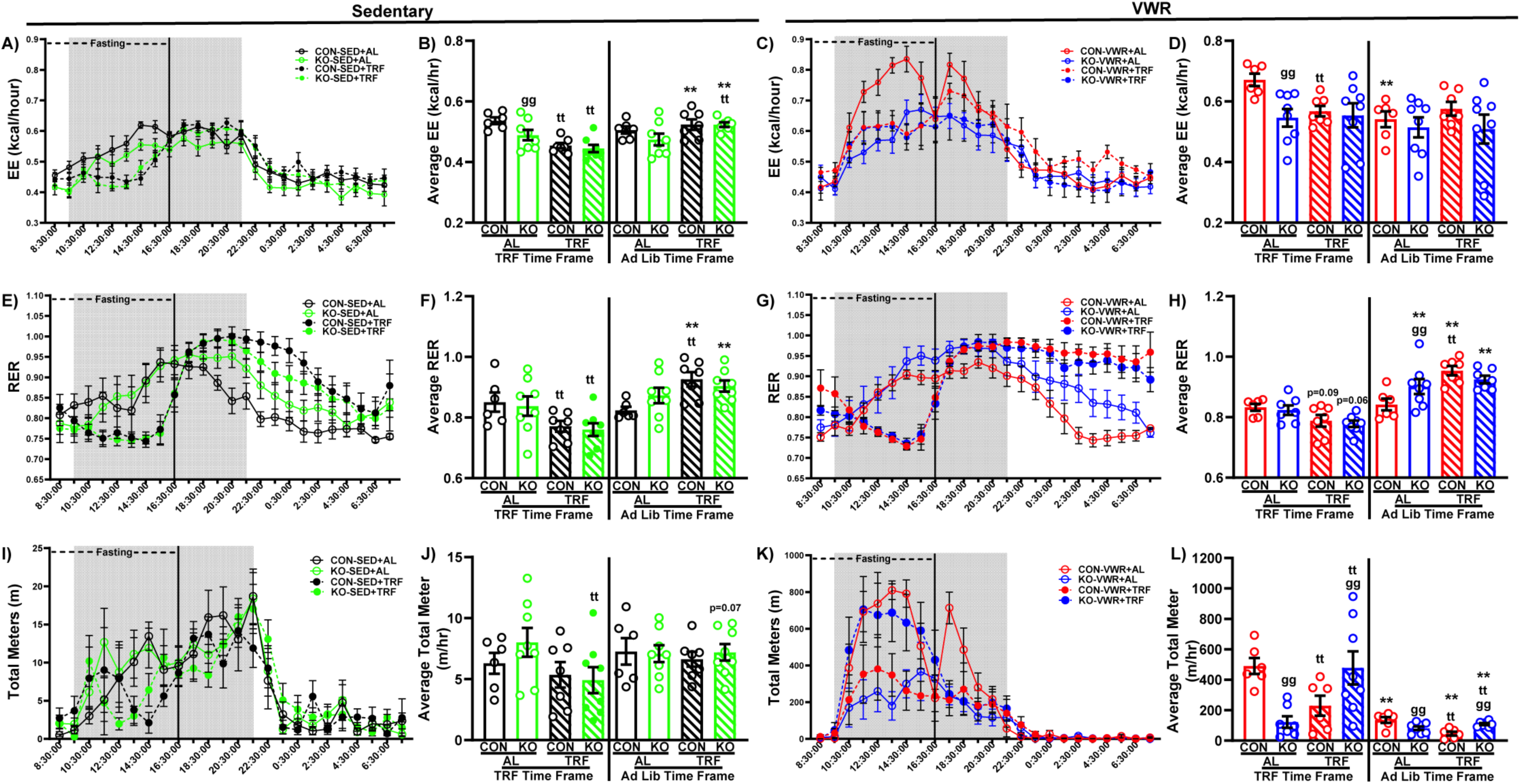
Hourly EE, RER and movement (left side – sedentary; right side – VWR). (A) Hourly EE for SED group; (B) Average EE for SED group in TRF Time frame vs Ad Lib time frame; (C) Hourly EE for VWR group; (D) Average EE for VWR group in TRF Time frame vs Ad Lib time frame; (E) Hourly RER for SED group; (F) Average RER for SED group in TRF Time frame vs Ad Lib time frame; (G) Hourly RER for VWR group; (H) Average RER for VWR group in TRF Time frame vs Ad Lib time frame; (I) Hourly cage activity for SED group; (J) Average movement for SED group in TRF Time frame vs Ad Lib time frame; (K) Hourly cage activity for VWR group; (L). Average movement for VWR group in TRF Time frame vs Ad Lib time frame; Data presented as mean +/- SEM (n=6-8 per group). Gg: p<0.05 genotype within feeding group and time frame, tt: p<0.05 TRF within genotype and time frame, ** p<0.05 time frame within genotype and feeding group.

**Figure 3.**
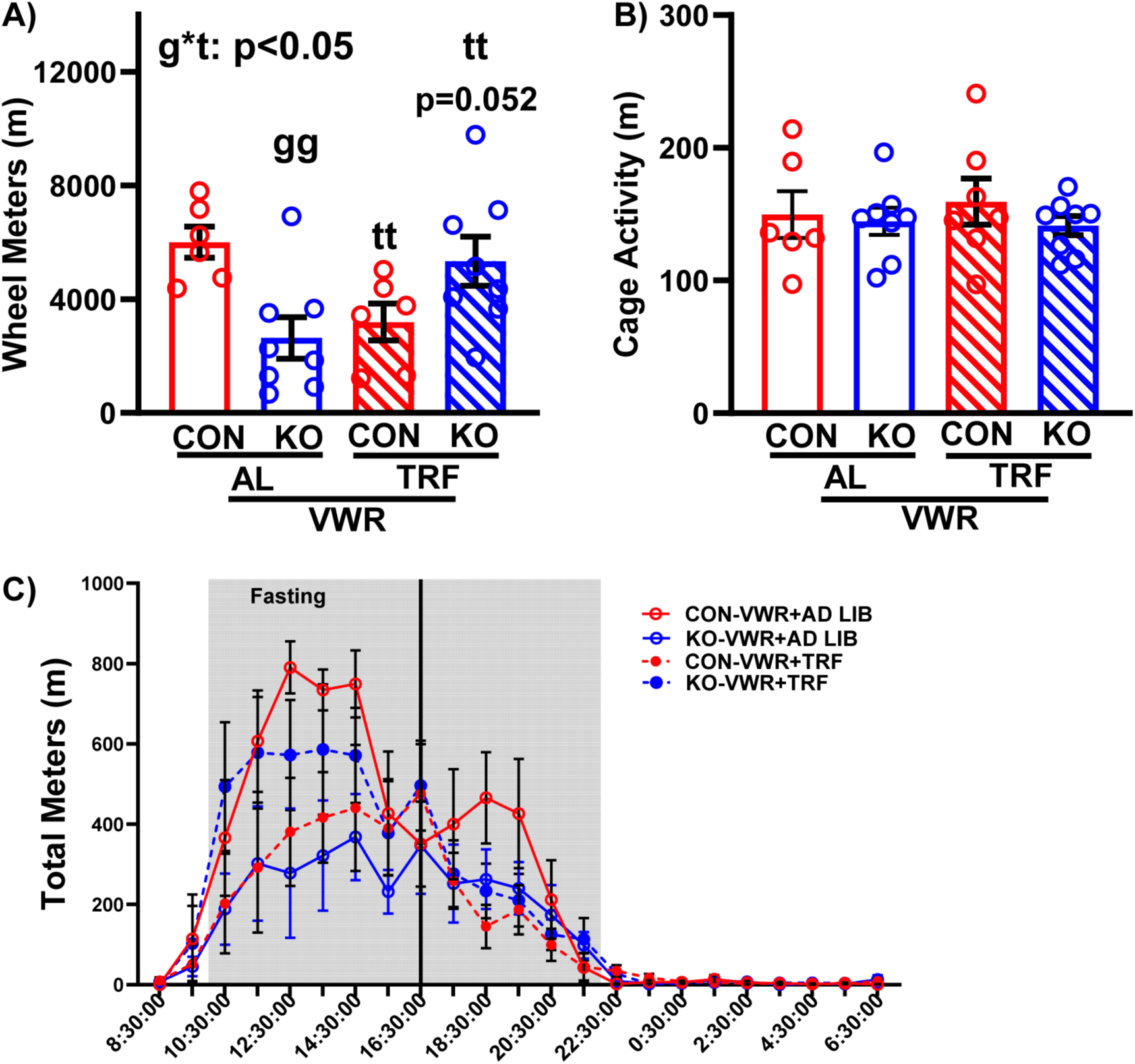
VWR group wheel meters and ambulatory cage activity (not including wheels) in indirect chambers. (A) Wheel running distance during data collection. (B) Cage activity distance during data collection with wheel running distance removed. (C) Day 3 of indirect calorimetry chamber data demonstrating similar behavior as day 4 (depicted in Figure 2K). g* t: genotype*TRF interaction p<0.05, v p<0.05 main effect of VWR, gg: p<0.05 genotype within feeding and activity, tt: p<0.05 feeding within genotype and activity.

### Hourly EE, RER, and movement

After identifying differences in Total Meters, RER, and EE, we investigated whether these factors exhibit temporal variations (hour by hour) between genotypes and treatments within either Sedentary or VWR conditions. While no significant differences in 24-hour EE were found among SED groups (Figure 1A), CON and KO SED+TRF mice both exhibited reduced EE during the TRF period (Figure 2B, p<0.05) and increased EE when they had access to food (p<0.05) compared to their within-genotype AD LIB counterparts. Additionally, KO SED AD LIB mice had lower EE than CON during the restricted feeding period (Figure 2B, p<0.05), and SED KO TRF mice showed higher EE compared to KO AD LIB during ad lib food access (p<0.05). In VWR groups, CON AD LIB mice displayed higher EE than both CON TRF and KO AD LIB during the food-restricted period (Figure 2D, p<0.05). Unlike the SED groups, no differences in EE were identified during periods when mice had access to food between VWR groups. However, CON AD LIB mice had decreased EE during ad lib food access compared to the food restriction period (p<0.05). Similar to the EE data in SED mice, RER measurements in SED groups were influenced by food availability when analyzed on an hourly basis. SED RER was lower in both TRF groups during restricted food access (Figure 2F, p<0.05) and higher during food access periods (p<0.05) compared to AD LIB groups. Furthermore, SED CON TRF mice exhibited higher RER during ad lib food access than CON AD LIB (p<0.05). In VWR groups (Figure 2H), TRF tended to lower RER in both CON (p=0.09) and KO (p=0.06) compared to AD LIB during food restriction (Figure 2H). During ad lib feeding, RER was elevated in both KO groups and CON TRF compared to the respective food restriction periods (p<0.05). KO AD LIB mice and CON TRF mice showed higher RER than CON AD LIB during the ad lib feeding period (p<0.05). Interestingly, RER was increased in VWR KO AL mice over VWR CON AL mice during the ad lib feeding period an effect not found in SED mice (Figure 2H). Similar to the 24-hour cage activity data, SED Total Meters was lower in KO TRF mice compared to AD LIB during food restriction (Figure 2J, p<0.05) and tended to be higher during ad lib feeding in this group compared to restricted feeding (p=0.07). The distinctive interaction observed in VWR mice regarding Total Meters in the averaged 24-hour cage activity analysis was further emphasized in the hourly analysis (Figure 2K-L). Specifically, KO AD LIB and CON TRF mice had lower Total Meters than CON AD LIB during restricted food access (p<0.05), while KO TRF Total Meters was higher than both KO AD LIB and CON TRF in the same period (p<0.05). Although all groups except KO AD LIB exhibited lower Total Meters during ad lib feeding compared to restricted feeding (p<0.05), the same pattern was observed across groups during ad lib feeding (p<0.05). To confirm that differences in average movement were not artifacts of single-day analysis, we examined the movement data for the VWR groups on the final day of habituation, before the primary measurements were taken (day 3), to determine if these traits were consistent. Indeed, all groups exhibited the same behavioral effects on day 3 as they did on day 4, as indicated by Total Meters (Figure 3C). These findings demonstrate that hepatic ketogenic insufficiency and TRF interact to create temporal differences in EE (Figure 2B and 2D), Total Meters (Figure 2L), and wheel running exercise (Figure 3A), especially during the normal dark cycle feeding period.

### HMGCS2 KO mice and skeletal muscle proteome adaptation to exercise

Skeletal muscle is a crucial site of energy metabolism during exercise. Since KO showed decreased non-resting EE during activity, we performed untargeted proteomics on skeletal muscle (whole gastrocnemius) to explore pathways that might influence EE. Our rationale was that skeletal muscle is a key site of energy metabolism during exercise and also a major site for ketone utilization. Driven by the calorimetry outcomes, conditions focused on SED+AL, VWR+AL, and VWR+TRF, all in CON and KO cohorts. The top upregulated and downregulated pathways are shown in Figure 4, using IPA analysis (IPA, cutoff Z-score = ±2.0, -log *p*-value > 1.3). Within genotype, condition-dependent comparisons are shown in Supplemental Figure 3. In the SED+AL condition, KO muscle shows a general reduction in proteins involved in oxidative phosphorylation (OXPHOS) (Figure 4A, Z-score= -4.92) compared to controls. VWR lessens this difference between KO and controls (Figure 4B), changing the Z-score from -5 to -3.25 for OXPHOS and from -4.92 to -2.25 for ETS, although they remain significantly suppressed in KO mice. When VWR is combined with TRF, the OXPHOS pathway is no longer a significantly diminished pathway in KO skeletal muscle (Figure 4C), and indeed the VWR+TRF condition in KO increases OXPHOS-related protein expression (Supplemental Figure 3). Pathway analysis of KO versus CON was combined to illustrate how VWR and VWR+TRF influence the gastrocnemius proteome (Figure 4D). In the VWR+TRF groups, KO shows higher protein levels linked to OXPHOS (Figure 4D, Z-score=2.77) than controls. Due to clear differences in pathways involving OXPHOS proteins across genotypes and TRF conditions, we cross-referenced these proteins with MitoCarta3.0 to identify known mitochondrial proteins within our dataset. The comparison between CON and KO mice revealed no differences in total mitochondrial proteins between KO and CON (Figure 4E), indicating that overall mitochondrial content differences do not cause variations in the OXPHOS pathway. We then normalized all OXPHOS proteins to total mitochondrial protein content. In the SED+AL condition, KO mice have significantly fewer OXPHOS proteins in skeletal muscle than controls, but VWR mitigates this difference regardless of whether it occurs under AL or TRF conditions (Figure 4F). Interestingly, VWR+TRF has contrasting effects across genotypes. In CON, VWR+TRF reduces OXPHOS protein levels in skeletal muscle, whereas in KO mice, it significantly increases them (Figure 4F, genotype × VWR+TRF < 0.05). Aside from mitochondrial proteins, IPA analysis shows that KO skeletal muscle has increased expression of various other pathways in response to VWR+TRF compared to CON. For example, the EIF2AK4 (GCN2) pathway responds to VWR+TRF, with a Z-score shifting from -3.24 in SED+AL to 2.60 in VWR+TRF in KO mice (Figure 4D). Other pathways affected by VWR+TRF in KO include neutrophil extracellular trap signaling, eukaryotic translation elongation, and ribosomal quality control signaling (Figure 4D).

**Figure 4.**
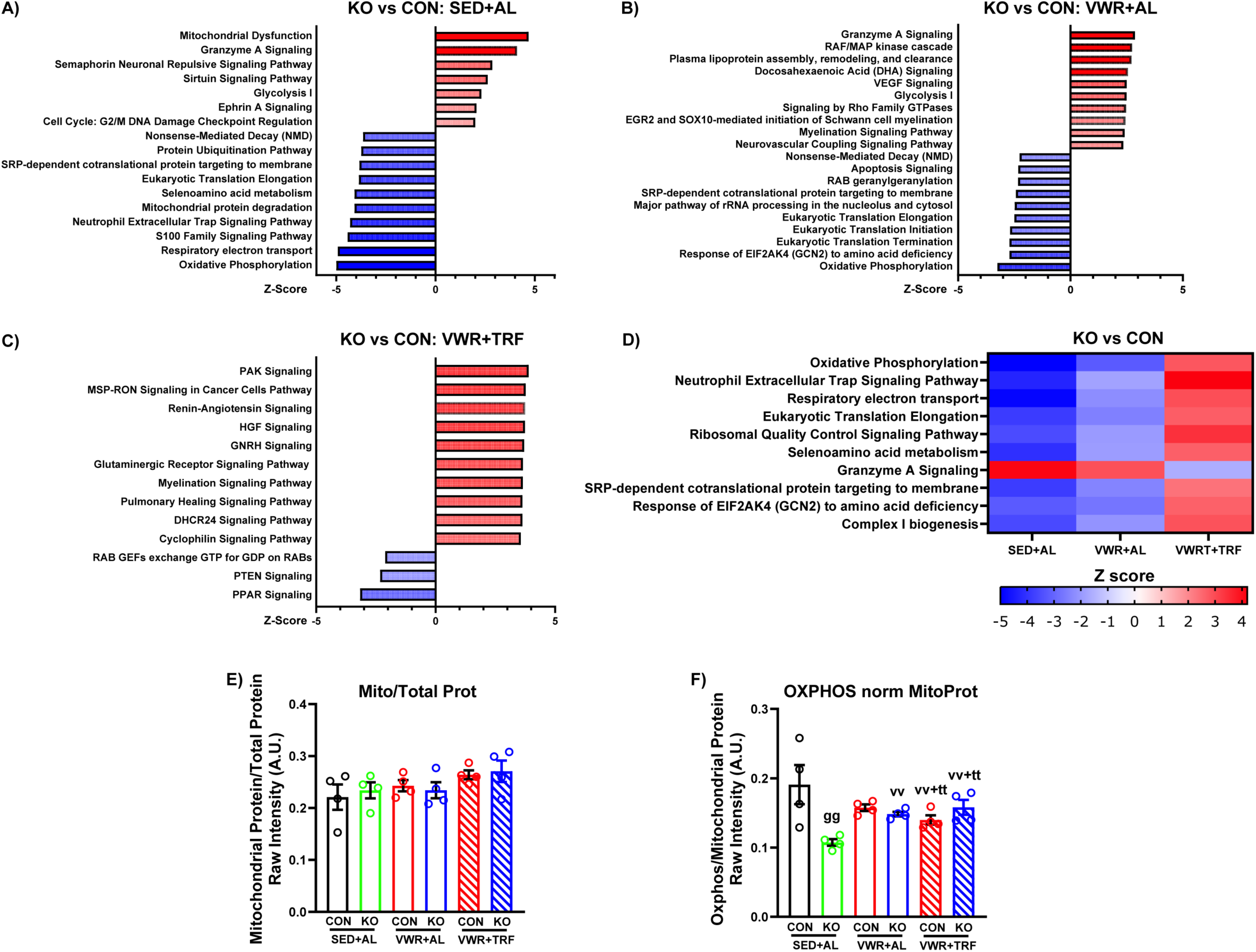
Comparison of adaptation between KO and control. (A) Pathway analysis for SED+AD LIB group KO over CON, (B) Pathway analysis for VWR+AD LIB group KO over CON, (C) Pathway analysis for VWR+TRF between KO over CON, (D)Heat map visualization for proteomic analysis across three different groups, (E) Mitochondrial protein over total protein expression across all groups. (F) OXPHOS protein normalized by mitochondrial protein. For panels (E) and (F), data presented as mean +/- SEM (n=4 per group). Gg: p<0.05 genotype within feeding and activity, vv: p<0.05 activity within genotype and feeding, and vv+tt: p<0.05 feeding and activity within genotype.

### Significant genotype difference in protein expression associated with glycolysis

While ketone metabolism pathways in muscle were not significantly different between genotypes (absolute Z-score < 2.0), we specifically examined proteins associated with KB metabolism due to its relevance to our study (Supplemental Figure 2). Z-scores were shown under all three conditions (Supplemental Figure 2A). Under SED+AL condition, CON has higher HMGCLL1 protein expression, a protein involved in leucine degradation and KBs synthesis (24). VWR+AL and VWR+TRF both decreased HMGCL1 protein expression in the CON but did not affect the KO (Supplemental Figure 2B, interaction genotype*VWR < 0.05; interaction genotype*VWR+TRF < 0.05). To our surprise, the genotype difference in protein expressions related to ketone utilization was minimal. Different interventions have minimal impacts on protein expression associated with ketolysis (Supplemental Figure 2C, 2D, & 2E). Since we observed a consistent increase in glycolysis-associated protein expression (Figure 4A and B), we examined the abundances of specific glycolytic enzymatic mediators. Gastrocnemius muscle from HMGCS2 KO showed lower HK2 protein expression across all three conditions (Figure 5), but higher levels of GPI and TPI1 (Figure 5; main effects of genotype, p < 0.05.) Under both AD LIB and TRF conditions, VWR increased GPI expression compared to SED+AL (Figure 5). When VWR was combined with TRF, KO had higher GPI expression than when VWR was combined with AL, but this increase was not seen in CON mice (Figure 5, genotype x TRF < 0.05). This genotype x TRF interaction was also captured in PFKM (catalyzes the rate-limiting glycolytic reaction) and TPI1 proteins (Figure 5). VWR plus TRF increased expression for PFKM, TPI1, and GAPDH in the KO compared to SED+AL, but this increase did not occur in CON muscle (genotype*VWR+TRF < 0.05). VWR+TRF increased PGK1 and PGM1 protein levels regardless of genotype compared to SED+AL (main effect p < 0.05). Across all conditions, KO generally showed higher PGM1 protein levels than CON. We also performed separate analyses within the CON and KO groups to visualize how different interventions affected skeletal muscle protein expression differently (Supplemental Figure 3). These results further demonstrate that liver HMGCS2 KO leads to substantially different regulation of multiple pathways in skeletal muscle in response to various interventions, highlighting the pronounced influence of hepatic KB production on skeletal muscle metabolism and mitochondrial profiles.

**Figure 5.**
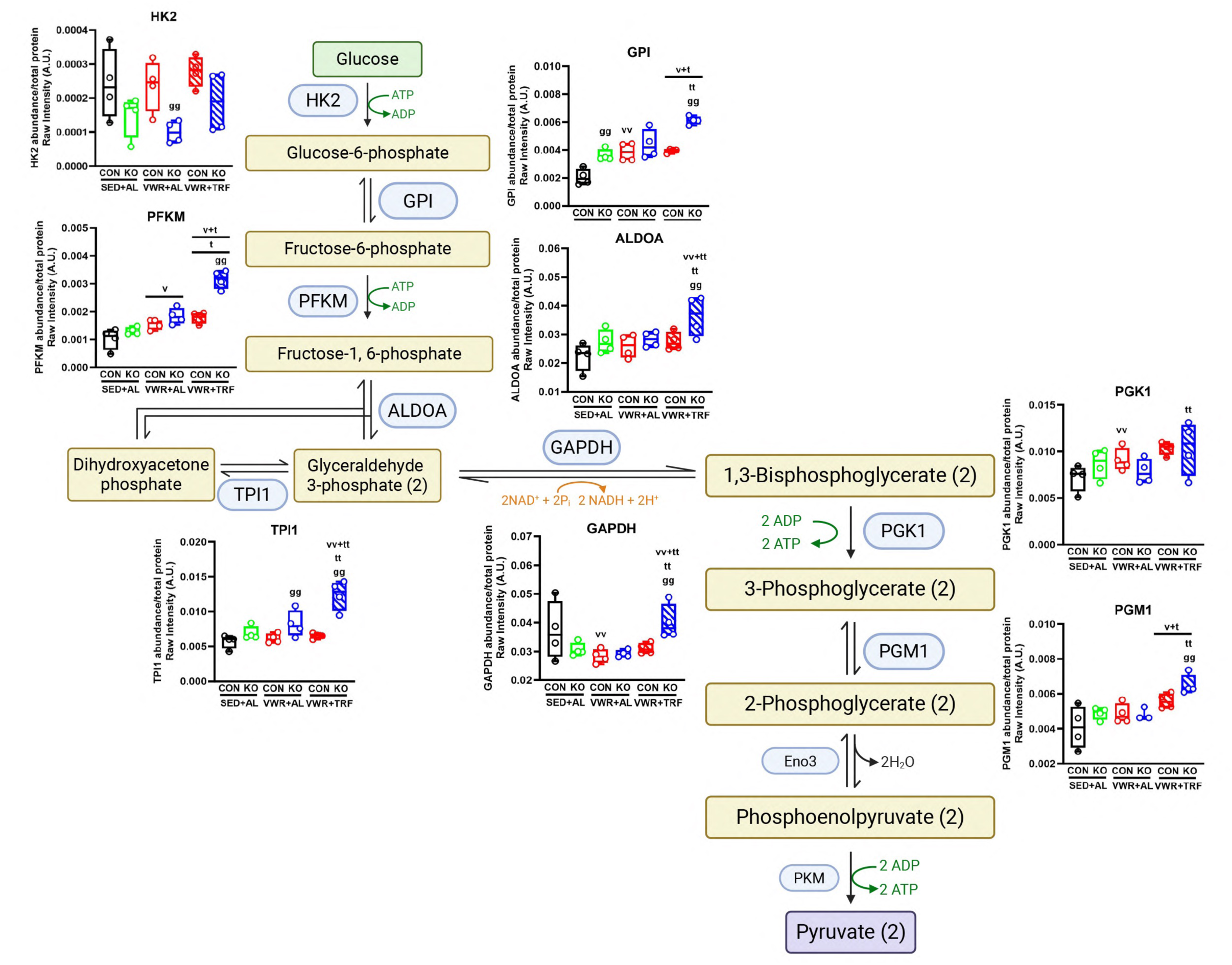
Significant genotype differences in glycolytic-associated proteins. Raw intensity protein abundance for glycolysis proteins: (A) HK2, (B) GPI, (C) PFKM, (D) TPI1, (E) GAPDH, (F) PGK1, (G) PGM1. Data presented as mean +/- SEM (n=4 per group). Gg: p<0.05 genotype within feeding and activity, vv: p<0.05 activity within genotype and feeding, tt: p<0.05 TRF within genotype and activity, and vv+tt: p<0.05 feeding and activity within genotype.

## DISCUSSION

Our main finding is that mice with liver-specific knockout of HMGCS2, which causes hepatic ketogenic insufficiency, show impairment in both total and non-resting energy expenditure in response to exercise compared to controls. This phenotype was latent in the sedentary state but was provoked by chronic exposure to daily voluntary wheel running. Additionally, the lower energy expenditure phenotype in knockout mice occurs in those with either daily ad libitum feeding (AL) or daily food restriction (TRF), which were combined with exercise to stimulate ketone production during chronic intervention. Other notable whole-body metabolic changes in the knockout mice include variations in movement efficiency, measured as energy expenditure per unit of movement, during both fasting and fed states, compared to controls. The ability of TRF to recruit augmented ketogenesis, abrogated in the KO, may influence movement frequency and efficiency. We also investigated muscle proteomics because muscle is a primary site for ketone utilization and makes a significant contribution to exercise-induced energy expenditure. Our results show that, under the SED+AL condition, liver ketogenic insufficiency markedly affects the muscle proteome by reducing OXPHOS proteins, but exercise through VWR with time-restricted feeding normalizes these responses relative to controls. Overall, our findings suggest that hepatic ketogenesis plays a crucial role in regulating both whole-body energy metabolism and skeletal muscle proteome adaptations to exercise and TRF.

As an alternative fuel source, ketone metabolism interacts with various energy pathways such as β-oxidation of fatty acids, de novo lipogenesis, sterol biosynthesis, and glucose metabolism (25). Because ketone metabolism is interconnected with multiple metabolic pathways, many studies have been conducted to clarify how it affects energy expenditure and substrate utilization. It is important to understand the biological effects of ketone metabolism on energy expenditure and to explore the potential impact of ketogenic strategies on energy balance. In this study, we show that hepatic ketogenesis influences energy expenditure under both fasting and exercise conditions. Compared to the SED group, VWR increased TEE, but this effect was almost abolished in the KO regardless of AL or TRF conditions. TEE consists of two main components: non-resting EE and resting EE. Further analysis revealed that only non-resting EE was affected by genotype, while resting EE was not. This aligns with the EE phenotype only occurring in KO mice undergoing VWR and not in sedentary mice.

Other notable whole-body phenotypes also appeared in the KO mice that are connected to the EE phenotype. During time in the indirect calorimetry cages, the KO ran less than CON in the AD LIB groups, suggesting that a lower amount of exercise may partially contribute to the lower non-resting EE phenotype in KO mice. However, this was not the case in the VWR+TRF treatment, where KO ran more than CON. Therefore, despite running more, KO still showed lower non-resting EE than CON. VWR increases energy expenditure, which in turn raises energy demand. During fasting, the liver boosts fatty acid oxidation to produce KBs and meet the demands of an upregulation in whole-body energy metabolism. When ketogenesis is impaired, VWR no longer increases non-resting EE, indicating that KB is a fuel that influences whole-body energy metabolism during exercise. What remains unknown is whether this effect on EE is driven by the synthesis of KB during exercise or the use of KBs in extra-hepatic tissues.

It is well known that ketogenesis mainly occurs in the liver, where fatty acids are broken down into acetyl-CoA and then converted into ketone bodies (KBs) through a series of enzymatic reactions. Besides terminal oxidation in the TCA cycle and very low-density lipoprotein secretion, ketogenesis functions as a primary pathway supporting fat disposal in the liver, contributing to the rate of hepatic fat oxidation, which is believed to protect against hepatic steatosis (26, 27). Hepatocytes use KBs to support fatty acid biosynthesis through both de novo lipogenesis (DNL) and polyunsaturated fatty acid (PUFA) elongation (12). Recent studies suggest that ketogenesis aids fatty acid biosynthesis by providing hepatocytes with acetyl-CoA, and the loss of HMGCS2 hampers fatty acid elongation and increases liver triacylglycerol (12). Fatty acid elongation can be energy-intensive, and ketogenesis may enhance energy expenditure by supporting this process. Ketone utilization can also impact overall body energy metabolism. Compared to fatty acid oxidation, ketone bodies are more energy-efficient, producing more ATP per molecule of oxygen consumed (P/O ratio) (28, 29). The impaired ketone body catabolism in knockout mice may contribute to this interesting phenotype.

KBs also exert many signaling functions that regulate exercise-induced adaptation in skeletal muscle. We investigated if there were specific skeletal muscle adaptations that could contribute to the lower TEE /Non resting EE phenotype seen in the liver HMGCS KO. Utilizing pathway analysis, we identified that the Sed KO group possess lower mitochondrial respiratory proteins than the equivalent CON mice, but this genotype difference was attenuated with VWR. Oxidative phosphorylation is significantly more efficient than glycolysis, and this higher OXPHOS protein expression is observed in KO, which may facilitate a more efficient energy production process and result in lower TEE and NREE. Because we did not measure mitochondrial respiratory capacities in skeletal muscle, we are unable to make conclusions about the efficiency of OXPHOS capacity between the CON and KO. Interestingly, we did not observe any significant differences in protein associated with ketolysis across the genotype. However, we did notice a significant difference in protein expression associated with glycolysis pathway: KO has higher glycolysis protein expression than CON. It suggested that in response to the lack of ketone as a fuel substrate, KO switches to glycolysis as one their major energy source. Previously studies have showed that ketogenic insufficiencies increase glycogen storage and glycogenolysis within liver It is plausible that ketogenic insufficiencies and the subsequent reduction in ketone availability alter muscle glycogen storage. This change in muscle glycogen storage could, in turn, induce distinct glycolysis protein expression patterns when comparing control (CON) and knockout (KO) groups. Again, one limitation is that we did not perform any function test and examine how the muscle utilizes ketone or glucose as their substrates, Therefore, we are unable to draw any conclusions regarding the efficiency of ketone metabolism or glycolysis between the CON and KO.

Skeletal muscle is also an energy consumer that detects nutrient availability which could possibly affect energy metabolism. Proteomic analysis revealed that KO had more significant changes in protein associated with general control nonderepressible 2 (GCN2) response to amino acid deficiency. GCN2 is a serine/threonine-protein kinase that senses amino acid deficiencies and plays a vital role in modulating amino acid metabolism in response to nutrient deprivation (30). GCN2 also promotes the phosphorylation of eIF2a, which facilitates adaptation to nutrient stress (30) and lowers protein synthesis via conserving energy and nutrients (31). While many studies have shown amino acid deprivation activates GCN2 (32, 33), little is known how ketone deficiency alters the GCN2 pathway. Proteomics analysis showed that skeletal muscle in the HMGCS2KO upregulates protein expression associated with GCN2 pathway, which as a result, could potentially trigger a series of changes to lower energy expenditure and conserve nutrients/energy. However, whether or not GCN2 pathway plays a primary regulatory role is unknown.

Our study has several limitations. We only performed this study using male mice. Previous studies suggest that sex differences in lipid metabolism exist with females demonstrating a higher fat utilization during exercise (34). We would anticipate that female mice would present a different response to the 16-week intervention under the various conditions. Sample size may have also been a limitation for outcomes, including 3-way ANOVA. Finally, we only performed proteome analysis of the gastrocnemius skeletal muscle and did not add functional measures of mitochondrial respiratory capacity, or glucose/fat oxidation,

In summary, this study utilizes liver-specific HMGCS2 knockout mice to demonstrate that a significant reduction in hepatic ketogenic capacity blunts exercise-induced energy expenditure and significantly alters mitochondrial and metabolic proteomic signatures in skeletal muscle. These baseline differences in skeletal muscle proteome between CON and KO then also likely effected responsiveness for various pathways to adapt to VWR, TRF, and TRF+VWR. Future investigation should focus on whether it is the metabolic pathways that synthesize KB within the liver or reduced utilization of KB in peripheral tissues like skeletal muscle that alters the capacity for exercise to modulate energy expenditure at the whole-body level.

---

**Supplement Figure 1.**
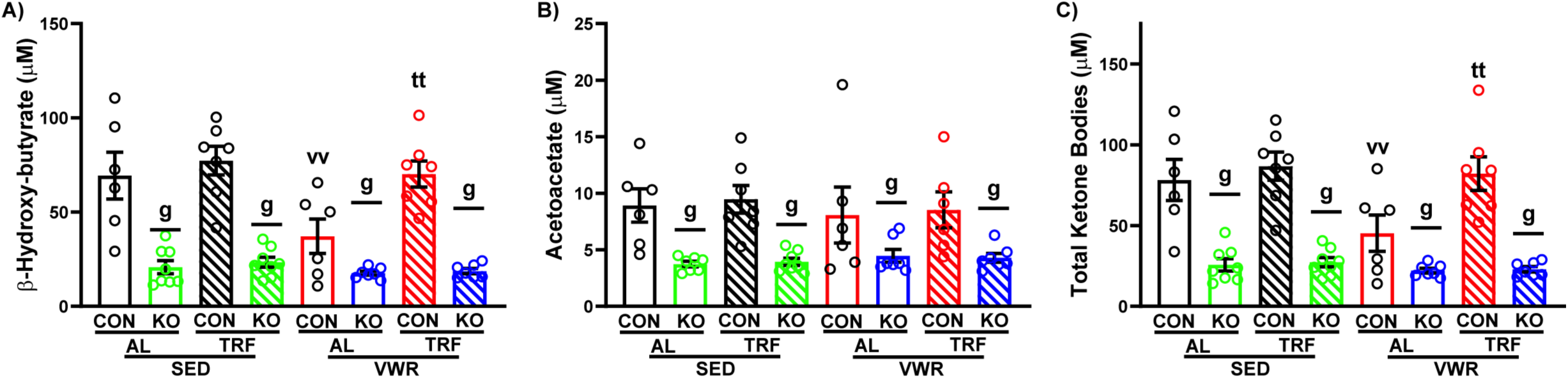
Serum ketone body levels. (A) beta-hydroxybutyrate serum levels (mM), (B) acetoacetate serum levels (mM), and (C) total ketone body serum levels (mM). Data presented as mean +/- SEM (n=6-8 per group). g: p<0.05 main effect of genotype, vv: p<0.05 activity within genotype and feeding, and tt: p<0.05 TRF within genotype and time frame.

**Supplement Figure 2.**
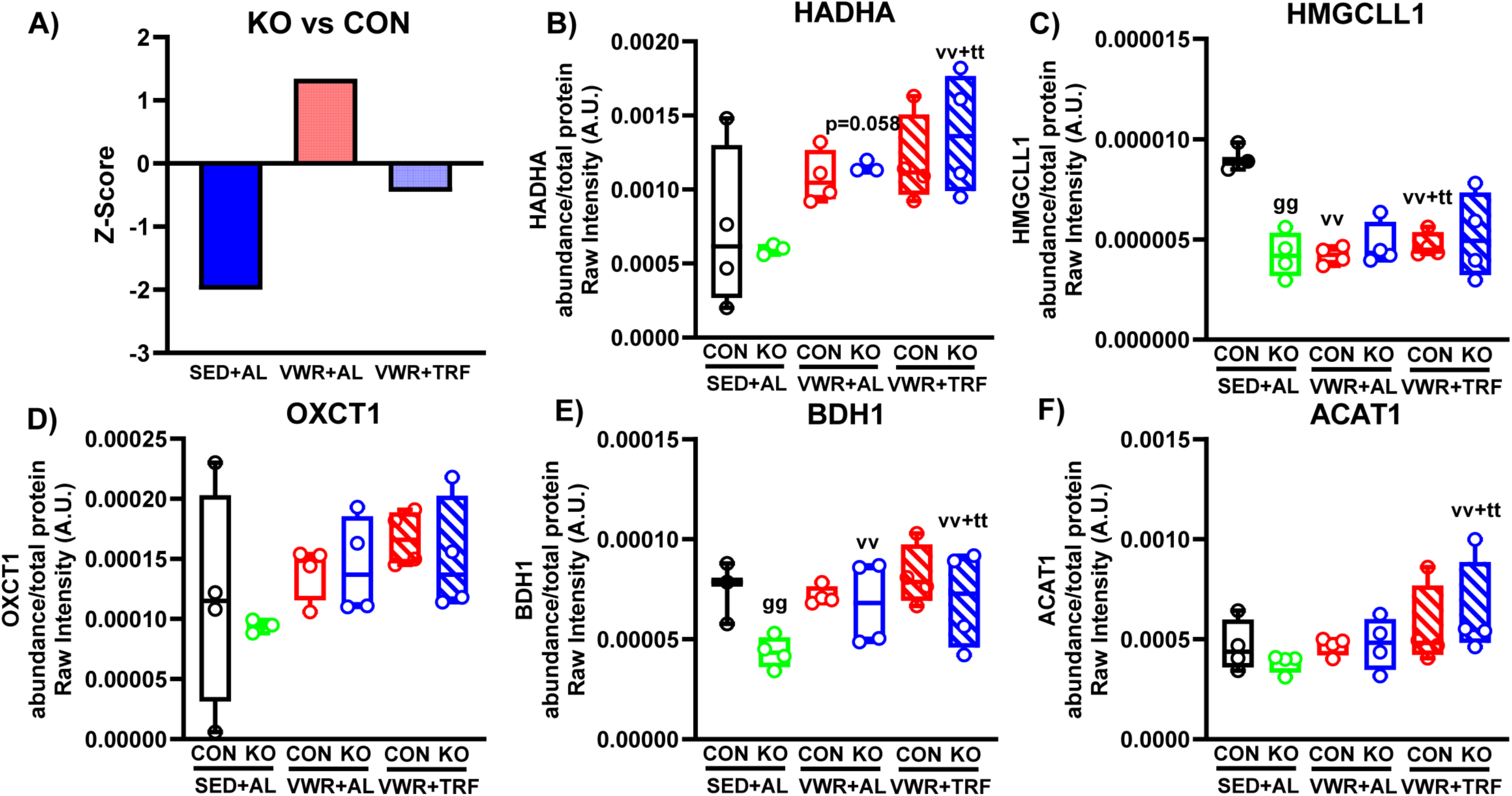
Minimal genotype difference in the ketone-metabolism-associated protein. (A) Z score across three conditions, (B) HMGCLL1 (C) BDH1, (D) OXCT1, (E) ACAT1. Data presented as mean +/- SEM (n=4 per group). gg: p<0.05 genotype within feeding and activity, vv: p<0.05 activity within genotype and feeding, and vv+tt: p<0.05 feeding and activity within genotype.

**Supplement Figure 3.**
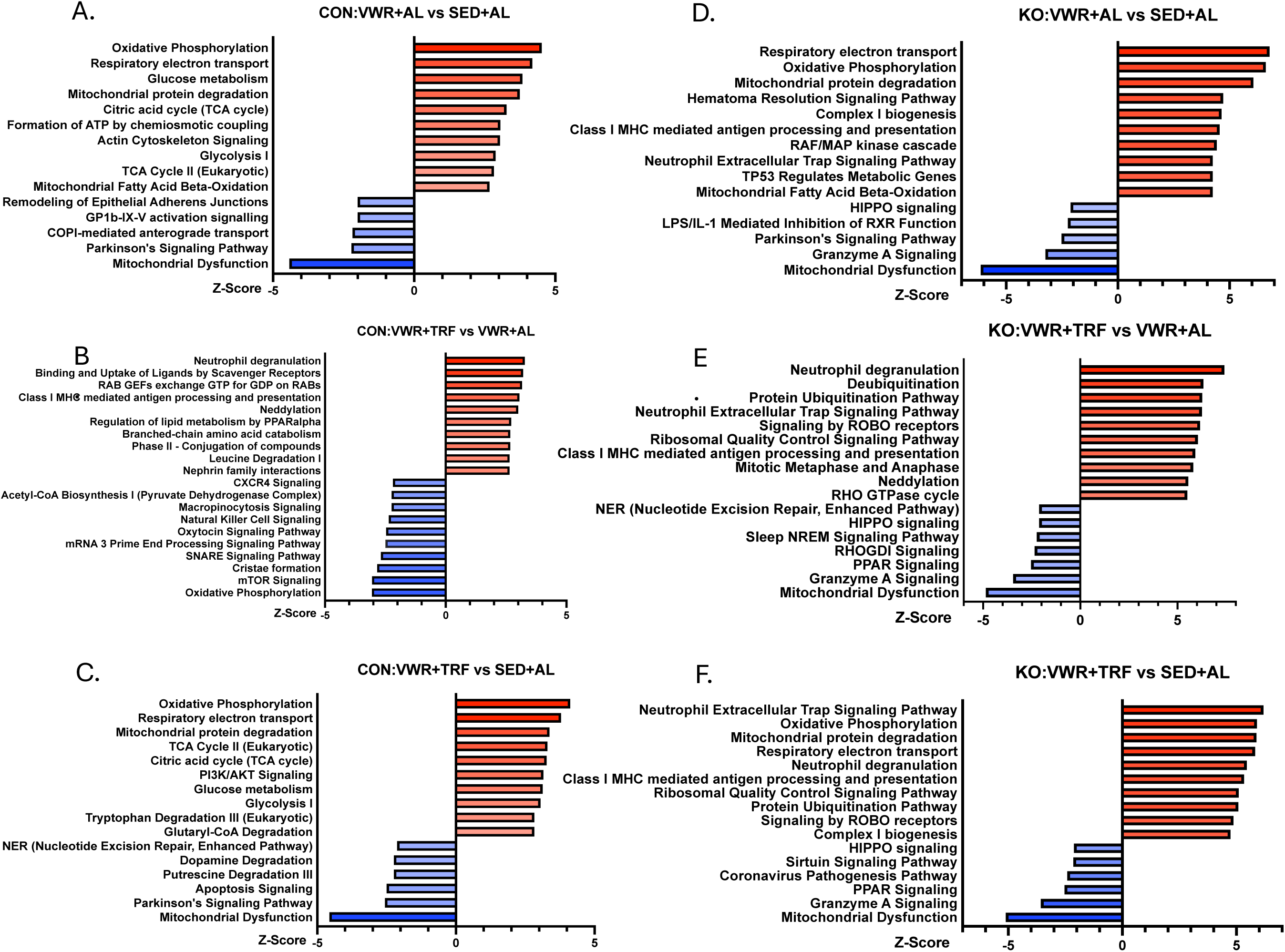
Comparison of adaptation within control and KO. (A) Pathway analysis for CON:VWR+AD LIB VS. SED+AD LIB, (B) Pathway analysis for CON: VWR+TRF VS. VWR+AD LIB, (C) Pathway analysis for CON: VWR+TRF vs. SED+AD LIB, (D) Pathway analysis for KO:VWR+AD LIB VS. SED+AD LIB, (E) Pathway analysis for KO: VWR+TRF VS. VWR+AD LIB, (F) Pathway analysis for KO: VWR+TRF vs. SED+AD LIB (n=4 per group).

